# Phage composition of a fermented milk and colostrum product assessed by microbiome array; putative role of open reading frames in reference to cell signaling and neurological development

**DOI:** 10.1101/714154

**Authors:** Stefania Pacini, Marco Ruggiero

## Abstract

Bacteriophages (phages), Earth’s most numerous biological entities, are natural constituents of alimentary matrices; in this study we describe the characterization of phage populations in a product obtained by fermentation of bovine milk and colostrum. Such characterizations were achieved using a microarray consisting of a chip covered in short DNA sequences that are specific to certain target organisms for a total of approximately 12,000 species. The only viruses evidenced by the array belonged to Siphoviridae, the largest phage family that targets bacteria and archea. The array yielded 27 iterations corresponding to a unique target. We discuss the putative role of some open reading frames of these phages in conferring health-supporting properties with particular reference to cells signaling and neurological development. We also describe the *in vitro* interaction of this fermented product with alpha-N-acetylgalactosaminidase, an enzyme whose activity in serum is elevated in neurodevelopmental disorders.

## Introduction

It is well assessed that consumption of probiotics under the form of fermented milk products such as yogurts or kefirs is associated with health benefits and, in particular, with improvement of immune system function (1, 2). It appears that most healthy properties are associated with the process of fermentation rather than with the microbes themselves. The same microbial strains that proved effective in improving the immune status of people living with HIV/AIDS, when administered as a probiotic yogurt (1), were ineffective when administered as pills containing only the encapsulated microbes and not the products of fermentation (3). According to the Authors of the latter study, “The inefficacy of the probiotic strains in preserving the immune function may be the result of using encapsulated probiotics versus the use of probiotic yogurt with the same probiotic strains in previous studies …” (3). It can be hypothesized that microbial metabolism, during the process of fermentation, leads to the production of molecules endowed with certain healthy properties. These molecules derive from inactive precursors present in the milk before fermentation. Such an hypothesis is corroborated by evidence demonstrating that in kefir, the process of fermentation leads to production of 236 unique, newly formed peptides from the microbial metabolism of bovine milk caseins (4). Many of the newly formed peptides show health-promoting properties due to their antithrombotic, immune-modulating, antimicrobial, antioxidant, mineral binding, and opiomimetic characteristics and to their effectiveness as ACE inhibitors (4).

In addition to peptides derived from casein metabolism, in 2011 we hypothesized that beta-galactosidase and sialidase produced by lactic acid bacteria and Bifidobacteria (5, 6) were responsible for metabolism of vitamin D-binding protein (DBP) leading to production of DBP-derived macrophage activating factor (DBP-MAF) (7), a molecule endowed with a number of properties (8, 9). Consistent with this hypothesis, consumption of a product deriving from fermentation of milk and colostrum, the latter being naturally rich in DBP, for three weeks, was associated with significant improvement of immune system function (7). Based on this experience, this product of fermentation of milk and colostrum containing probiotic strains typical of yogurts and kefirs, has been used in a nutritional approach for chronic conditions since 2014 (10, 11, 12). In this paper, we describe how the DBP-MAF naturally formed during fermentation of this product interacts with human alpha-N-acetylgalactosaminidase (nagalase) and how such an interaction compares with that of purified DBP-MAF.

In addition, we subjected this product to detailed genetic analysis using the Axiom Microbiome Array, a tool that is described as “the next generation microarray for high-throughput pathogen and microbiome analysis” (13). According to the Authors from Lawrence Livermore National Laboratory, Thermo Fisher Scientific, and Kansas State University, “The array contains probes designed to detect more than 12,000 species of viruses, bacteria, fungi, protozoa and archaea, yielding the most comprehensive microbial detection platform built to date” (13). The sensitivity of the array is such that it is able to detect pathogenic bacteria such as Shigella and Aspergillus at 100 genome copies, and viruses such as vaccinia virus DNA at 1,000 genome copies. Using this novel method to better characterize the microbial composition of the fermented product quoted above, we discovered that its biodiversity was much greater than previously anticipated, and we further documented the presence of bacteriophages (phages) belonging to the family of Siphoviridae. Such a presence did not come as a surprise since phages are the planet’s most numerous biological entities and are naturally present in practically all alimentary matrices from American dairy products (14) to sweet and salted Italian baked products (15) and even to mother’s breast milk (16).

In recent years, interpretation of the role of phages changed from that of “enemies” to that of “allies” in a shift of perception reminiscent of that occurred for bacteria that went from “bugs” to be eliminated at all costs, to constituents of the healthy human microbiota (17, 18). Here we describe the phage composition of this probiotic product, and discuss the putative role of the open reading frames (ORF) of some phages in conferring health supporting properties.

## Materials and Methods

The Axiom Microbiome Array was performed by Eurofins Microbiology Laboratories Inc. (New Berlin Wisconsin, USA) on a commercially available fermented product designated “Freeze-Dried Bravo - Colostrum and Probiotic Complex” (Silver Spring Sagl, Arzo-Mendrisio, Switzerland). This product derives from 48 hrs of fermentation of bovine milk and colostrum at room temperature. After 48 hrs, the product undergoes the process of freeze drying. The nutritional characteristics of the product are described in Antonucci et al. (2019) (19). The array used for this analysis is a chip covered in short DNA sequences that are specific to certain target organisms (a total of approximately 12,000 species are included).

For each target organism, the Microbiome Detection Analysis Software (MiDAS; Thermo Fisher) looks at the ratio of probes that successfully bound to their target (Probes Observed) to the total number that could potentially bind (Probes Expected) and uses this ratio to calculate the Initial Score, which is proportional to the likelihood of the target existing in the tested sample. According to Thissen et al. (13) “The initial score is the log likelihood ratio for the target being present in the sample if no other targets are present, *vs* no targets being present in the sample. This value gives information on what the maximum possible contribution of that target is to the holistic model of the sample, based on the probes observed when interrogating a sample with the Axiom Microbiome Array.

The conditional score gives an indicator of the actual contribution of each target to the model of the sample; it is the log likelihood for a model including the target *vs* a model without the target. As the conditional score takes into account the presence of other targets, it can be lower than the initial score for a given target if there are probes in common between targets.” The MiDAS software is based on the Composite Likelihood Maximization Method (CLiMax) algorithm developed at Lawrence Livermore National Laboratory (13). Once the software identifies the target that it considers most likely to be present in the actual sample, the probes that correspond to the chosen target are removed and the process is repeated until adding new species no longer adds explanatory power to the software’s model of the composition of the sample. The products of this iterative analysis become and are defined as “Iterations.” Probes, with signal intensity above the 99^th^ percentile of the random control probe intensities and with more than 20% of target-specific probes detected, are considered positive. In other words, a target organism is considered present when the percentage of observed probes is higher than 20% of the expected probes. In the example by Thissen et al. (13) vaccinia virus was detectable when there were 1,000 or more copies of the virus DNA in the tested sample. The number of probes observed at 1,000 copies was 78 out of 293 probes expected, or about 26%, and this value was considered positive for the presence of the virus. This assay method enables some sort of quantification; in the example quoted above, the number of probes observed at 10,000 copies of vaccinia virus was 148 out of 286 probes expected, or about 52%.

Study of *in vitro* interaction between the fermented product and human nagalase was performed by R.E.D. Laboratories (Zellik, Belgium) where human nagalase and DBP-MAF were purified. The experiments were performed using microtiter plates coated with a specific antibody able to capture human nagalase. Samples were incubated with a standardized dilution of a pool of 300 human sera from healthy subjects and, after 1 h incubation and exhaustive washing, complexes formed by nagalase and purified DBP-MAF, used as positive control, or nagalase and the fermented product, were detected with a horse radish peroxidase conjugate of a rabbit antibody. In order to establish the kinetics of interaction between nagalase and DBP-MAF or nagalase and the fermented product, the serum pool was mixed either with 200 ng of purified DBP-MAF, or with two dilutions (1:10 and 1:100) of the fermented product in phosphate buffered saline (PBS). The mixture was then incubated at room temperature for 4, 24, 48, 72 and 120 h. Values for nagalase binding activity in the absence of DBP-MAF or the fermented product, with only PBS in the reaction mixture, were taken as 1.00. The experiment was repeated twice and the results reported in Fig. 1 are the means of the two experiments.

**Fig. 1.**
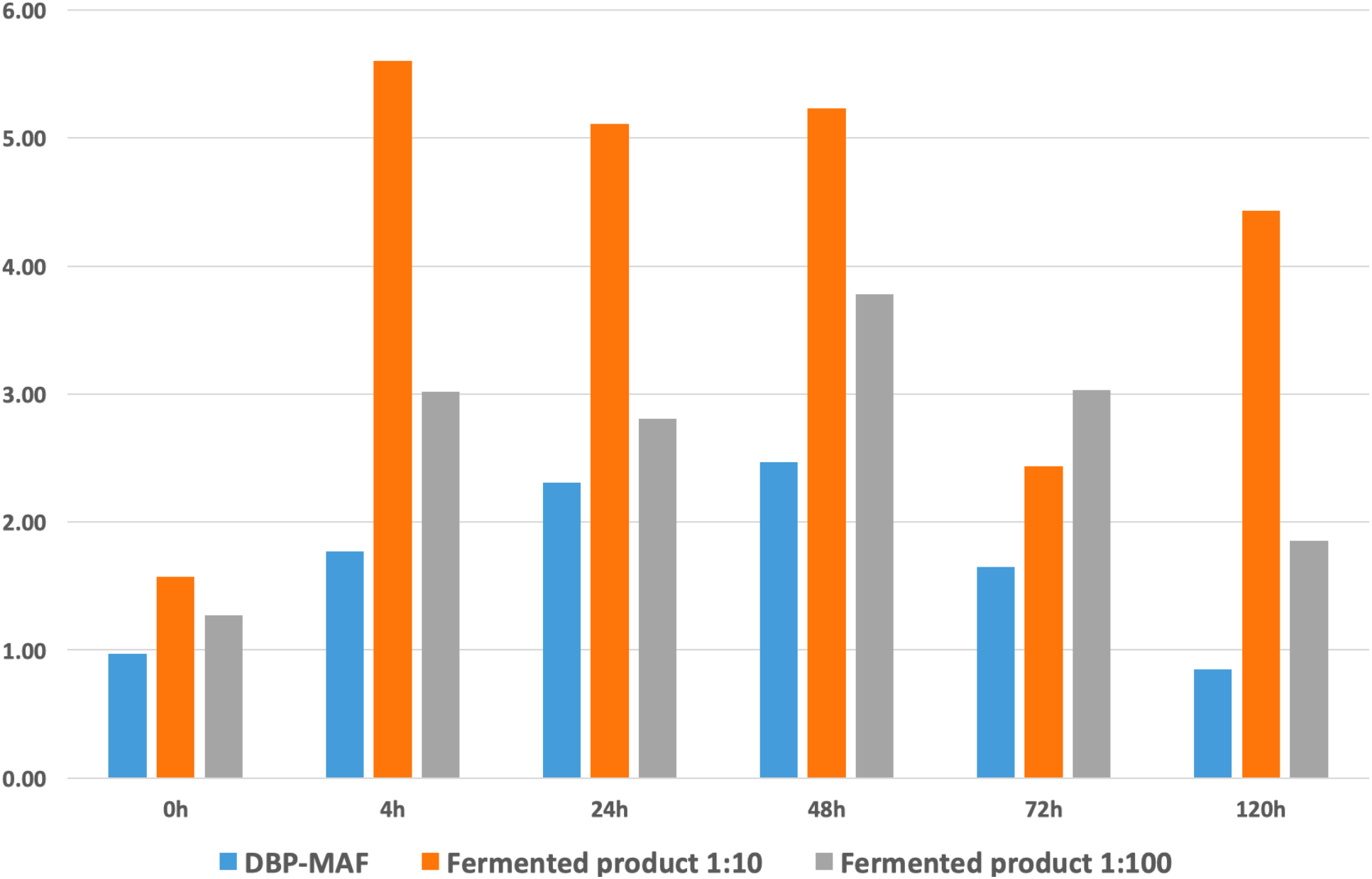
Binding activity against human nagalase.

## Results

### Phage composition

The Axiom Microbiome Array did not evidence in the analyzed product any pathogenic microbes considering the sensitivity of the method as described in Thissen et al. (13). This result provides significant information on the safety of the product. This observation is also consistent with the results of the microbiological analyses for contaminants that are routinely carried out for the product in the context of the company’s quality control system. The array demonstrated the presence of a very high number of targets corresponding to probiotic species. The species evidenced by the array belong to the families of Streptococcaceae, Lactobacillaceae, Leuconostocaceae, Bifidobacteriaceae, and Thermaceae. Detailed description of the prokaryotic targets identified by the array will be presented in another paper. Here, we focus our attention on double-stranded DNA viruses belonging to Siphoviridae family, the only type of viruses evidenced in the product; these represent the largest phage family that targets bacteria and archea (20).

Table 1 shows the phage composition of the product as far as phages are concerned. The array provided 27 iterations corresponding to unique Siphoviridae targets. The phages are ordered in descending sequence with those showing the highest percentage of observed probes at the top of the list. Although the assay is not strictly quantitative, based on the results of Thissen et al. (13), it can be argued that higher percentages corresponded to higher number of viral copies.

**TABLE 1.**
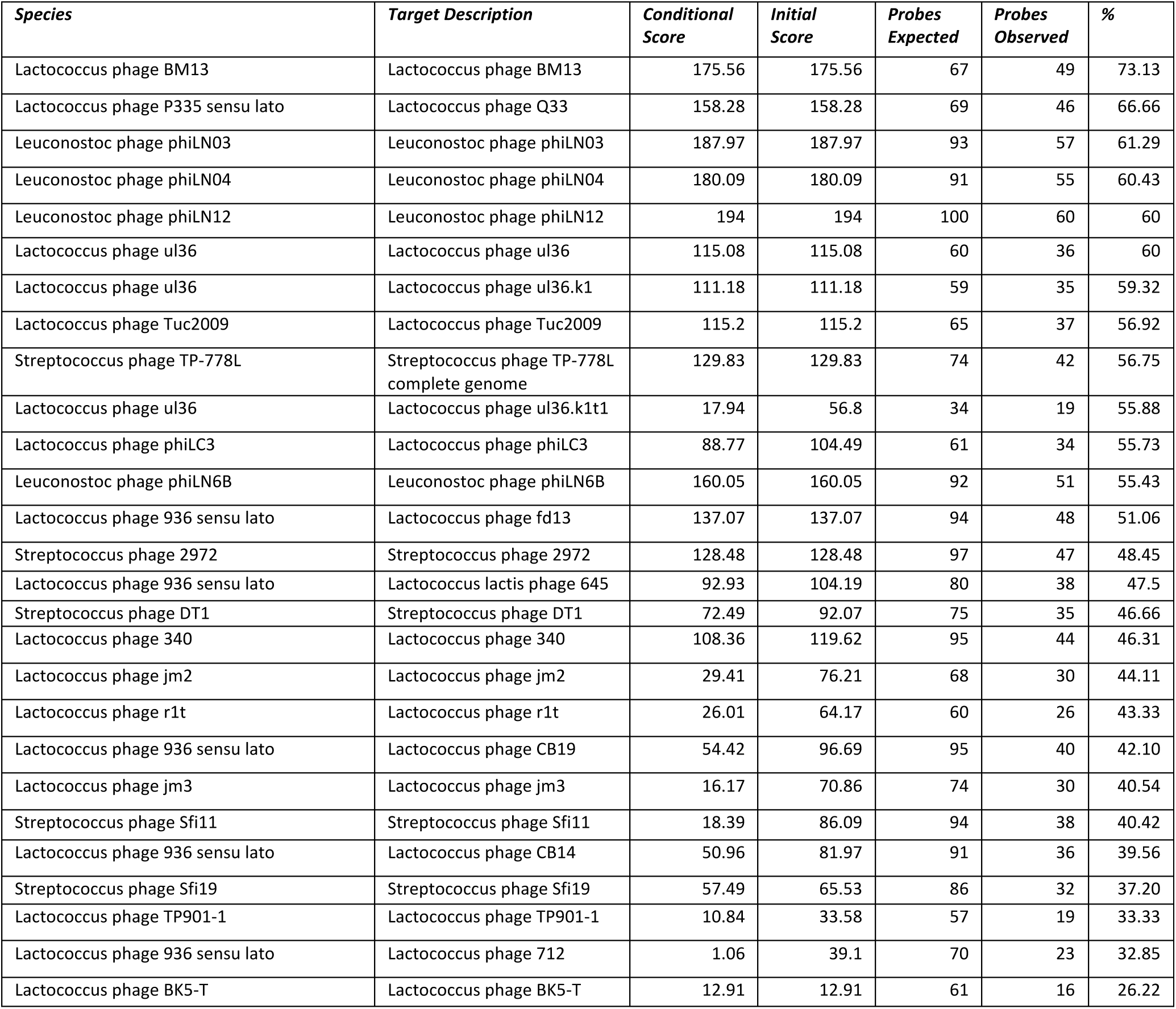

### In vitro interaction between the fermented product and nagalase

As shown in Fig. 1, purified DBP-MAF, used as positive control, bound human nagalase only after 4 h incubation, reached a peak at 48 h, and returned below baseline values at 120 h. The fermented product had an initial value (at time 0) higher than that observed with PBS alone, thus demonstrating immediate, intrinsic DBP-MAF activity against human nagalase. At every time point, both dilutions of the fermented product showed significantly higher activity in comparison with purified DBP-MAF. At 120 h, the activity of both dilutions of the fermented product was still well above baseline, whereas DBP-MAF did not show any residual activity. Since the fermented product was diluted 10 and 100 fold, it may be argued that its activity against human nagalase is more than 100 fold higher than that of purified DBP-MAF. It is worth considering that these results were obtained *in vitro*, that is in the absence of any variable or confounding factor that may hamper interpretation of results previously observed in clinical settings (12).

## Discussion

Phages have been used for human consumption for more than 100 years (21). In 2019, the Bacteriophage for Gastrointestinal Health (PHAGE) study evaluated the safety and tolerability of supplemental phage consumption. This randomized, double-blind, placebo-controlled crossover investigation performed by Authors from prestigious US research institutions, concluded that a mixture of phages comprising Siphoviridae, the family of phages described in the present study, was safe in humans (22).

Here we describe the main features of some of the phages evidenced by the Axiom Microbiome Array in the product object of this study as they potentially relate to human health.

Lactococcus phages BM13 and Q33, the most represented in the analyzed product, have a genome showing close relatedness to phages infecting Enterococci and Clostridia, a feature that is not shared by other lactococcal P335 phages (23). The ability to infect and induce lysis of Enterococci and Clostridia may bear significant consequences for human health. Enterococci are opportunistic pathogens capable of acquiring resistance to antibiotics and have emerged as important healthcare-associated pathogens (24). A number of Clostridia is associated with human disease. For example, alterations of the gut microbiota associated with antibiotic treatment stimulate the growth of resistant strains and the onset of Clostridium difficile infection (25). Therefore, phages able to infect and eliminate Enterococci and Clostridia could be useful in the management of conditions associated with these pathogens, and in particular, in the presence of antibiotic-resistant strains.

Lactococcus phage ul36 encodes a protein, Sak, that is homologous to a human recombination protein, RAD52, that plays a crucial role in DNA repair, genome stability and prevention of carcinogenesis (26, 27). Such an homology is present at the amino acid, phylogenic, functional, and structural levels and raises the possibility of a common origin with Sak being an ancestor of RAD52. A study by Authors from Canada, Switzerland and USA, demonstrated that purified Sak bound single-stranded DNA preferentially over double-stranded DNA, and promoted the renaturation of long complementary single-stranded DNAs in the context of DNA repair. Interestingly, Sak was able to self-assemble even in the absence of DNA with the formation of toroidal structures (26). This feature of Sak is interesting in the context of DNA repair, in particular, if it is compared to the self-assembly of a major protein contributing to human genome stability, the tumor suppressor protein p53, “the guardian of the - *human* - genome” (28). It was demonstrated that four p53 molecules need DNA to self-assemble on two half-sites of the polynucleotide to form a tetramer that consists of a dimer of dimers whose stabilization is assured by protein-protein and DNA base-stacking interactions (29). It appears that the ability of Sak to protect genome stability is more “primordial” as the information for self-assembly is all contained in the protein structure without the need for interactions with damaged DNA. This feature may provide Sak with the advantage of being less restricted in its ability to repair DNA and guarantee genome stability. Self-assembly of Sak in toroidal structures raises the interesting possibility of non-chemical signaling between the protein and DNA, possibly through biological quantum entanglement between the electron clouds of toroids of DNA and the electron clouds on the surface of toroidal Sak (for reference on biological quantum entanglement between electron clouds of nucleic acids, see 30). It is well known that DNA is organized in toroid units within sperm chromatin and in the head of phages (31, 32) and condensation of a constrained DNA molecule into a toroid increases its tension and modifies the electromagnetic properties associated with the distribution of electrical charges on its surface (33). It could then be hypothesized that toroidal structures of DNA and proteins such as Sak may become entangled because of their spatial geometry; this phenomenon would not be dissimilar from the quantum entanglement at the level of the protein tubulin in neuron microtubules that is thought to be at the basis of human consciousness (34). It would also be consistent with the recent paper by Authors from The Netherlands entitled “Is the Fabric of Reality Guided by a Semi-Harmonic, Toroidal Background Field?” (35). It is interesting to notice that DNA itself is endowed with an intrinsic degree of consciousness (36) that obviously does not rely upon quantum entanglement at the level of tubulin. It is also interesting to notice that ancient viruses may have been responsible for the onset of consciousness in humans (37) and, therefore, it is at least theoretically plausible that the intrinsic consciousness of the DNA of viruses and humans may have become entangled at different levels.

Although the concept of quantum entanglement may appear esoteric and distant from the daily routine of biological entities, quantum coherence and entanglement are at work in phenomena such as the origin of life (38), photosynthesis, a process at the basis of today’s life on this planet (39), biology of the retina (40), bird navigation in the skies (41), fish communication in the water (42), and human consciousness (for reference in addition to Hameroff et al. (34), see a recent study using Magnetic Resonance Imaging of the human brain (43). It is therefore of interest to consider that each phenomenon of quantum entanglement is associated with formation of a wormhole through the geometry of entanglement (44), a phenomenon that could explain the apparent reversal of direction of time in the experiments of Mothersill et al., a reversal of time described by the Authors as “temporally displaced awareness after the fish become entangled” (42).

Going back to the role of Sak in the context of DNA repair, it is worth noticing that Sak is able to bind RecA and stimulate homologous recombination reactions. Not surprisingly, human RAD52, an evolutionary derivative of Sak, was recently defined “a novel player in DNA repair in cancer and immunodeficiency” and was attributed a tumor suppressor function (45). These functions of Sak/RAD52 are consistent with a recent clinical observation in a case of multiple myeloma (19). It is also worth noticing that RAD52 was demonstrated to suppress HIV-1 infection in a manner independent of its ability to repair DNA double strand breaks through homologous recombination (46). Although there are no direct evidences for a role of Sak protein in suppressing retroviral infections, since this activity resides in the DNA-binding and self-interaction domains of RAD52 that are homologous to Sak, it is tempting to speculate that Sak of phage ul36 may also have anti-retroviral activities by binding to retroviral cDNA and suppressing integration of the infecting virus (46). We are not aware if the probiotic yogurt described by Irvine et al. in 2010 (1) contained phages and, in particular, Lactococcus phage ul36; if this were the case, however, the significant positive effects on people living with HIV/AIDS may be attributed also to the anti-retroviral properties of Sak.

Lactococcus phage TUC2009 shows interesting properties as far as detoxification of carcinogenic metals is concerned. It is well known that lactic acid bacteria such as those present in the examined product participate in detoxification of toxins that may contaminate food, examples of which are aflatoxin M1 (47), polycyclic aromatic hydrocarbons (PAHs), heterocyclic amines (HAs) and pthalic acid esters (PAEs) (48). It is also known that other strains present in the product, such as Streptococcus thermophilus, protect against toxicity of cadmium, a carcinogenic metal toxicant. Such a protection occurs by reducing the levels of cadmium in blood and ameliorating alterations in the levels of glutathione and malondialdehyde that are associated with cadmium toxicity (49). The presence of phage TUC2009 represents an interesting addition to the detoxification properties of the examined product as they were described in Blythe et al. (12). This is due to increased expression of the gene cadA that is about 20-fold higher in uninfected Lactococcus than in the bacteria infected by phage TUC2009 (50). This gene codes for a cadmium efflux ATPase and reduced expression following phage infection is associated with intracellular accumulation of cadmium. This effect of phage TUC2009 infection appears to be specific and not a generalized response to lytic phage infection. Thus, infection of Lactococcus by another lytic phage, c2, elicited an opposite response as far as expression of cadA was concerned resulting in increased transcription of the gene (50). It is tempting to speculate that bacteria infected with phage TUC2009 are more efficient in eliminating cadmium from the human organism as they are unable to expel it otherwise; cadmium would remain bound to bacterial proteins (51) and be eliminated from the organism together with the lysed bacteria.

Streptococcus phage TP-778L may be of interest in the context of the phenomenon designated viral superinfection exclusion. The gene ltp of this phage codes for a protein, Ltp, that is a lipoprotein attached to the outside of the cytoplasmic membrane. Ltp has the role of preventing viral superinfections by blocking injection of DNA of the infecting virus into the cytoplasm of the host cell (52). When the lipid-anchor domain is removed, the resulting truncated Ltp works as a broad-spectrum virus-resistance protein in the context of superinfection exclusion (53) and is able to protect the host cell against other, different, lactococcal phage species (54). Superinfection exclusion is mediated by interaction between negatively charged amino acids in the repeat domains of Ltp and the positively charged carboxyl terminus of the Tape Measure Protein (TMP) of superinfecting phages, as it was demonstrated for phage P008 (53). It is worth considering that a similar protein-protein interaction mediated by charged amino acids may be responsible for a conceptually analogous type of viral infection exclusion between Lactococcus phage fd13 and Epstein-Barr virus (EBV, see below). We are not aware of interactions between the negatively charged amino acids in the repeat domains of Ltp and the huge number of proteins that, having positively charged sequences, interact with heparin (55). However, considering that positively charged heparin-binding proteins comprise a number of extracellular proteins, growth factors, chemokines, cytokines, enzymes and lipoproteins, it is worth speculating that interaction between these and the exposed negatively charged sequences of phage proteins such as Ltp may have a role in regulating a variety of biological processes.

The complete genome sequence of Lactococcus phage phiLC3 is described in a paper published by Norwegian Authors in 2003 (56). In this paper, the Authors compare the genome of the phage to the genome of its relatives in Lactococci and Streptococci. Of interest is the sequence ATAAAAAATAGGAGAGTAAAATG of ORF88 that codes for a holin. Holins are proteins that form oligomers in the cytoplasmic membrane and lead to formation of holes that, together with the action of lysins, lead to cell death. The amino acid sequence coded for by ORF88 of phage phiLC3 shows significant (77%) identity with other holins, namely those of phages TUC2009, TP901-1, and a putative phage of Lactococcus casei. The percentage of identity was calculated over 31 amino acids. Such a significant percentage of identity leads to speculation of a common origin and it is interesting to note that holins perform essential functions also in eukaryotic cells. Two members of the eukaryotic Bcl-2 family, the multi-domain pro-apoptotic Bax and Bak proteins, display holin behavior that is not shared by other members of the Bcl-2 family (57). In eukaryotic cells, Bax and Bak exert their pro-apoptotic function in response to apoptotic stimuli by causing formation of holes in the mitochondrial outer membrane with consequent release of cytochrome c and other proteins into the cytosol to trigger the cascade of caspases (58). Since mitochondria evolutionarily derive from bacteria (59), it is not surprising that proteins able to form holes in bacteria exert the same action in mitochondria. Disruption of apoptosis is associated with the development of cancer, and Bax and Bak function as tumor suppressors thanks to their holin-like activities (60). It is known that phages induce tumor destruction and it was hypothesized that this anti-cancer activity of phages was mediated by interaction with the cells of the immune system, notably, the macrophages (61). Future research will elucidate whether holin-based mechanisms similar to that of tumor suppressors Bax and Bak may also be responsible for phage-induced cancer destruction.

Lactococcus phage fd13 has a receptor binding protein essential for host interaction that shows a significant degree of homology at the amino terminus with a number of other phages, namely bIL170, phi645, phi7, p272 (AF539443), P113G, sk1, jj50, p2, jw30, jw31, jw32, P008, phi936, phi712, and P475. Since all these phages have different host ranges, it is assumed that the amino terminus of the receptor binding protein is conserved and may serve the purpose of non-specific binding to the host cell (62). This leads to the interesting hypothesis of using phages to fight eukaryotic viral infections through protein-protein interactions as it was recently proposed for phage therapy of EBV infection (63). In the case of EBV, and possibly of other human herpes viruses, phages may prevent infectivity thanks to competition between proteins with KGD (Lys-Gly-Asp) motifs encoded by phages and herpes virus proteins required for membrane fusion and virus entry into eukaryotic cells. In the case of EBV, the KGD motifs involved in such a competition are those of in the gp24 head vertex protein of T4-like phages and those in the glycoprotein gHgL of EBV. Lactococcus phage fd13 shows a very conserved DGK sequence at the amino terminus (62) that may be considered “complementary” to KGD motifs of EBV and possibly of other human herpes viruses. The electrostatic interaction between amino acids with opposite charges (K and D) and the hydrophobic interaction between paired G may stabilize the binding of KGD to DGK motifs. Since KGD motifs of EBV mediate the infection of epithelial cells and B lymphocytes, it can be argued that DGK motifs of Lactococcus phages binding to KGD motifs may prevent infection by EBV and other herpes viruses.

Streptococcus phage 2972 shows a peculiar receptor binding protein that is coded for by ORF20. At the carboxyl terminus, this protein contains six collagen-like repeats motifs whose biological function is to provide elasticity and confer stability to the triple helix structure (64). It can be hypothesized that these motifs may interfere with the self-assembly of eukaryotic collagen monomers in fibrillary structures (65). Such a feature may prove interesting in the context multistep carcinogenesis (66) since collagen is actively involved in tumor progression as it constitutes the scaffold of tumor microenvironment and promotes tumor infiltration, angiogenesis, invasion and migration (67). Since phages ingested with a fermented product come in contact with the mucosa of the entire gastrointestinal tract, it may be hypothesized that such an effect may be of interest for prevention and treatment of malignant cell proliferation arising from the epithelium of the digestive tract. It cannot be ruled out, however, that such an effect may occur also at distant sites as discussed below.

The complete genomic sequence of Lactococcus phage DT1 was published in 1999 (68). The sequence of ORF32 that codes for a 26 kDa protein is of particular interest as it comprises an ATP–GTP binding motif A (P-loop). Phosphate-binding (P-loops) are found in ATP and GTP binding proteins (G-proteins). Their primary structure typically consists of a glycine-rich sequence followed by a conserved lysine and a serine or threonine (69). In molecular oncology, mutations in the context of P-loops bear important consequences as they are associated with oncogenes highly represented in human cancers such as the members of the ras gene family that code for signal transducing G-proteins in the same range of molecular weight of the protein coded for by ORF32. Mutants with three extra amino acids inserted in the P-loop of Ras proteins show significantly decreased affinity for GDP that leads to increased preference for GTP binding with increased oncogenic potential. In addition, in these mutants, GTP hydrolysis is drastically reduced thus leading to prolonged permanence of the oncogenic signal (70). About 33 years ago, we demonstrated that G-proteins coded for by oncogenic ras mutants, not only are associated with mitogenic signals that cannot be turned off, but also make the cells responsive to non-physiologic signals that, therefore, assume the role of tumor promoters. In particular, we demonstrated that cells transformed by transfection with the oncogene EJ/T24-H-ras, but not their normal counterpart, were able to couple a muscarinic receptor agonist, carbamylcholine, with calcium influx and polyphosphoinositide metabolism, all hallmarks of mitogenic signaling (71). Presence of a nucleotide-binding P-loop in the protein coded for by ORF32 of phage DT1 opens interesting perspectives in counteracting oncogenesis associated with mutated G-proteins. It can be envisaged that the phage protein acts as a decoy for GTP binding thus depriving the oncogenic protein of its substrate. It may be argued that mitogenic, signal-transducing G-proteins are localized inside cells as they are attached to the internal layer of the membrane through isoprenyl moieties, fatty acyl moieties, and electrostatic interactions (72), and they are therefore inaccessible to phage proteins. Such an argument however has been recently disproved by demonstration that phages are actually capable of penetrating into human cancer cells and remain active inside cells for at least 24h (73). Another argument against a role for phage proteins as tools against molecular carcinogenesis consists in the fact that ingested phages reside in the gastrointestinal tract and would therefore prove irrelevant for cancers arising in other parts of the body. However, this argument was also disproved by observation that in animals and humans, phages translocate from the gut to other organs (74); this phenomenon, coupled with the ability of phages to penetrate into eukaryotic cells, is probably responsible for the continuous and dynamic virus-prokaryote-eukaryote gene flow (73). As a matter of fact, ingested phages have been found in sera of healthy individuals since 1971 (75). Researchers from the Polish Academy of Sciences demonstrated since 1987 the presence of phages in blood and urine after oral administration of specific phage preparations aimed at treating bacterial infections resistant to antibiotics; no such phages were detected in those patients prior administration, thus confirming that the phages found in blood and urine came from the gut (76). Phages are able to reach blood through different routes of administration including the intranasal route (77). Once in blood, phages are fully active and able to kill pathogenic bacteria; in a recent paper, it was demonstrated that phage phiHP3 was able to infect, lyse and ultimately kill extra-intestinal pathogenic Escherichia coli (ExPEC), a leading cause of bloodstream infections, when present in human blood together with metals such as iron, magnesium and calcium (78).

Lactococcus lactis bacteriophage r1t was studied in detail as far as regulation of gene expression in responses to an anti-tumor chemotherapeutic agent, Mitomycin C, was concerned (79). The chromosomal region of interest of the phage comprises two genes, rro, that codes for the phage repressor, and tec. It was demonstrated that Mitomycin C, an agent that promotes the switch of r1t from the lysogenic to the lytic life cycle, increased 70-fold the expression of tec (79). These results may bear relevance in the context of counteracting the side effects of Mitomycin C chemotherapy. It is known that Mitomycin C leads to oxidative DNA damage and decreases global DNA methylation in a sex-specific pattern in mice; it is also known that most of the changes induced by chemotherapy in the pre-frontal cortex of female mice are similar to those occurring during the brain’s ageing (80). Considering the effects of Mitomycin C on phage r1t gene expression, it could be hypothesized that chemotherapy also alters the balance between phages and Lactococci in the microbiota, with consequences on the functions regulated by the microbiota and, in particular, on the immune system. Stimulation of lytic effects by Mitomycin C should then be taken into account, and strategies to support the Lactococci population could be implemented with the goal of counteracting the increased lethality of phages such as r1t.

Lactococcus phage CB19 belongs to a group of phylogenetically related heat-resistant phages that comprises P1532, SL4, CB13, and CB20. These phages show a remarkable resistance to heat and remain infectious after heat treatment at 80°C for 5 min (81). Molecular characterization of heat-resistant mutants of phage CB14, a related phage that infects the same host as CB19, suggests that resistance to heat is due to the amino acid sequence of TMP, a structural protein within the phage tail. The size of this protein in CB19 is 996 amino acids that is the same size of the other heat-resistant phages SL4, CB13, and CB20, whereas the size of the protein in heat-sensitive, wild-type CB14 is 999. The size of the heat-resistant mutants of CB14 is, however, 959 (81). These data seem to indicate that shorter TMPs are associated with resistance to heat. Such an interpretation is corroborated by evidence that phage P1532, heat-stable up to 90°C, shows an even shorter protein composed by 916 amino acids. TMPs of phages infecting Lactococcus strains have hydrophobic regions that assemble in transmembrane-spanning domains that serve different functions for phage infectivity including the recruitment of chaperones (82). These features, associated with heat-stability as it is the case for phage CB19, may represent an interesting starting point to harness the power of phage-derived, heat-stable proteins to interact with membrane-spanning receptors involved in mitogenic signal transduction. For example, phage-derived heat-stable proteins with hydrophobic domains could interact with receptors such as the Epidermal Growth Factor Receptor that is coupled with intracellular polyphosphoinositide mitogenic signaling as we demonstrated in 1988 (83). Thanks to the heat resistance of phage CB19, such an approach would be particularly indicated in combination with anti-cancer hyperthermia (84).

Streptococcus phages Sfi11 and Sfi19 are two phage types that, although related, are significantly different since they infect different strains of Streptococcus thermophilus and have different amino acid composition and tail morphology with Sfi11 showing thinner tails in comparison with Sfi19 (85). Both phages show a number of homologies with Streptococcus thermophilus phage TP-J34. Of particular speculative interest for its implications in cancer is the homology between ORF35 of TP-J34 and gp53 of Sfi11 with 100% match identity (53). gp53 is a putative transcription regulator (86). Since a transcription regulator is, by definition, a protein able to interact with DNA, it is tempting to speculate that a transcription regulator may interact also with p53, either at the DNA or at the protein level. Indirect support for this hypothesis comes from observation that peptides isolated from phage display libraries are able to bind human and murine p53 (87). It can be speculated that interaction between gp53 and p53 may restore transcriptional activity of mutant p53; this could be a promising strategy to harness the potential of p53 in protecting cells from carcinogenesis through cell-cycle arrest, senescence, and apoptosis (88).

## Conclusions

Phages have been used in therapy of human diseases for more than hundred years, that is soon after their discovery in 1915, when it was enthusiastically thought that they could cure almost any disease (21, 89). Although phage therapy is only now being rediscovered in Western countries (66), it is worth noticing that pharmacies in the country of Georgia and in the Russian Federation have been selling commercial phage cocktails for decades as they belong to the armamentarium of common practitioners (90). In the pre-antibiotic era, phages were conceived as means to kill pathogenic bacteria according to the principle of “My enemy’s enemy is my friend” (91), but now it is accepted that they may be helpful in a number of diseases ranging from cancer (92) to autism (93) as they are able to bypass anatomical and physiological barriers and have been found in human body’s compartments that were previously considered purely sterile (94). It is also worth considering that many of the biological effects of phages are mediated by their interactions with the immune system. For example, it has been demonstrated since 2009 that tumor-specific phages induce tumor destruction through activation of Tumor-Associated Macrophages as they promote a switch from a tumor-promoting M2-polarized microenvironment to a more M1-polarized milieu (61). This feature of phages is of particular interest when immunotherapies for cancers involving macrophage stimulation are considered (11). Thus, stimulation of macrophages in tumors in the absence of tools that direct them toward an M1 phenotype may lead to counterproductive effects since Tumor-Associated Macrophages of the M2 phenotype are known to promote tumor proliferation and to be associated with a poor prognosis in numerous cancers (95).

Although phage therapy has never been discontinued in Eastern Europe, in the Western world phage therapy faces regulatory obstacles that limit its diffusion. For example, the Food and Drug Administration of the USA classifies phages within the category of “biologics,” a category that comprises anything derived from biology and not from chemical synthesis such as, for example, blood products, antibodies, proteins or viruses that may prove efficient in treating specific diseases. Since biologics are more complex than chemically synthesized small molecule drugs, contain more than one single molecule, may have variable activity, may contain components exerting other functions, and have more complex pharmacokinetics, their regulation and approval are much more challenging (78). All these hindrances are well summarized by the Belgian Author Fauconnier who, in April 2019, wrote “After decades of disregard in the Western world, phage therapy is witnessing a return of interest. However, the pharmaceutical legislation that has since been implemented is basically designed for regulating industrially-made pharmaceuticals, devoid of any patient customization and intended for large-scale distribution. … The repeated appeal for a dedicated regulatory framework has not been heard by the European legislature, which, in this matter, features a strong resistance to change despite the precedent of the unhindered implementation of advanced therapy medicinal product (ATMPs) regulation. It is acknowledged that in many aspects, phage therapy medicinal products are quite unconventional pharmaceuticals, and likely this lack of conformity to the canonical model hampered the development of a suitable regulatory pathway. However, the regulatory approaches of countries where phage therapy traditions and practice have never been abandoned are now being revisited by some Western countries, opening new avenues for phage therapy regulation” (96).

These considerations, however, may not apply to phages already and constitutively present in alimentary matrices. We propose that phages naturally present in fermented products may yield additional benefits as compared to purified phage preparations (90) in a manner analogous to that observed for probiotic yogurts *vs* encapsulated probiotics (1, 3). Just like probiotics support the immune system in a much more efficient manner when in their environment, meaning fermented milk, phages may show the best health-supporting qualities when in their environment, that is, in the milieu where their hosts are present in great numbers. It is of interest to note that the presence of phages does not impede the process of fermentation when the biodiversity of the bacterial strains is elevated as is the case for the product described in this paper where the Axiom Microbiome Array evidenced hundreds of different bacterial strains, with the majority of them pertaining to the kefir grain component of the product. Observing the efficiency of fermentation, it is tempting to speculate that bacteria and phages have reached a point of equilibrium in the microflora of the kefir grains and are able to co-exist. It is also tempting to speculate that this naturally achieved equilibrium may favor a sort of entanglement with the human microbiota, a microbiota naturally rich in phages (97). Therefore, we feel that the answer to the question posed by the article in Science provocatively entitled “Does a sea of viruses inside our body help keep us healthy?” is definitely positive (98).

As far as the data on DBP-MAF and nagalase are concerned, this is the first time that such a direct interaction is demonstrated. Binding of DBP-MAF, or of the fermented product, to nagalase may help explaining the biological observations described in previous papers (9-12, 19, 99). The results reported here may bear particular relevance in the field of neurodevelopmental diseases where elevate serum nagalase activity is associated with clinical symptoms and successful immunotherapy is associated with decrease of such an activity (100). The 100 fold higher activity of the fermented product in comparison with purified DBP-MAF can be explained taking into consideration two elements.

1. Naturally formed DBP-MAF in the fermented product is associated with vitamin D_3_, fatty acids and glycosaminoglycans that are normal constituents of milk and colostrum (101). Since 2013, we proposed a molecular model describing how non-covalent association of these constituents in a multi-molecular complex significantly enhanced the biological activity of DBP-MAF (102). This model was independently confirmed by a recent study demonstrating that association with vitamin D_3_ significantly increased the immune stimulating activity of DBP-MAF (103).

2. Phages in the fermented product synthesize proteins with activities superimposable to that of DBP-MAF. For example, the protein RAD52 that is the human homolog of the protein Sak encoded by Lactococcus phage ul36, shows significant similarity with the active site of DBP-MAF. The sequences of human RAD52 (P43351) and vitamin D-binding protein (P02774) were obtained and aligned using the Align tool of Uniprot (uniprot.org). Alignment showed a striking similarity between the active site of DBP-MAF that is the peptide TPTELAK, and the peptide DPAQTSD of RAD52. Since it is well known that phages stimulate the immune system, and it is also known that such a stimulation is associated with macrophage activation (61), it is tempting to speculate that DBP-MAF-like proteins encoded by phages contribute to this effect by interacting with nagalase.

## Acknowledgements

The Authors wish to thank Dr. Jerry Blythe for critically reviewing the manuscript and suggesting changes to improve intelligibility. The Authors also wish to thank Dr. Tanja Mijatovic, PhD, Chief Scientific Officer at R.E.D. Laboratories, for insights on the significance of laboratory tests and for contributing to the understanding of the role of nagalase in health and disease.

## Authors’ Contributions

Stefania Pacini, MD, PhD and Marco Ruggiero, MD, PhD contributed equally to this paper.

## Disclosures

Marco Ruggiero is the founder and CEO of Silver Spring Sagl, the company producing the fermented product described in this paper. Stefania Pacini works as Quality Control Responsible Person for Silver Spring Sagl. Both had no prior knowledge of the results of the Axiom Microbiome Array that was independently performed by Eurofins Microbiology Laboratories Inc. (New Berlin Wisconsin, USA) on a commercially available lot as part of the company’s quality controls. Likewise, they had no prior knowledge of the results of the experiments with nagalase that were independently performed by R.E.D. Laboratories (Zellik, Belgium).

## Advisory

No information in this paper is intended or implied to be a substitute for professional medical advice, diagnosis or treatment.

## Notes

#### Summary of Updates

This version of the manuscript has been revised to add novel experimental data and refocus the paper on the significance of phages in neuroimmunology.

